# Brain-like functional specialization emerges spontaneously in deep neural networks

**DOI:** 10.1101/2021.07.05.451192

**Authors:** Katharina Dobs, Julio Martinez, Alexander J.E. Kell, Nancy Kanwisher

## Abstract

The last quarter century of cognitive neuroscience has revealed numerous cortical regions in humans with distinct, often highly specialized functions, from recognizing faces to understanding language to thinking about what other people are thinking. But it remains unclear why the cortex exhibits this high degree of functional specialization in the first place. Here, we consider the case of face perception, using artificial neural networks to test the hypothesis that functional segregation of face recognition in the brain reflects the computational requirements of the task. We find that networks trained on generic object recognition perform poorly on face recognition and vice versa, and further that networks optimized for both tasks spontaneously segregate themselves into separate systems for faces and objects. Thus, generic visual features that suffice for object recognition are apparently suboptimal for face recognition and vice versa. We then show functional segregation to varying degrees for other visual categories, revealing a widespread tendency for optimization (without built-in task-specific inductive biases) to lead to functional specialization in machines and, we conjecture, also brains.

## Introduction

Although controversial for centuries^1^, the idea of localization of function in the human brain is now supported by overwhelming evidence. Many regions of cortex are selectively activated by a specific perceptual or cognitive task, and when disrupted produce selective impairment of that same task^2–5^. But it remains unknown, and indeed largely unasked, *why* the brain exhibits this high degree of functional specialization. One possibility is that functional specialization in the cortex is an accident of evolution, which can more easily add modules to solve new problems, than redesign an entire system from scratch. Another possibility is that functional specialization allows mental processes to be selectively modulated, whether over short time scales (attention) or longer ones (development). A third (nonexclusive) possibility is that functional specialization in the brain arises for computational reasons, with distinct brain regions arising only for tasks that cannot be solved with more generic machinery. Here we test this third hypothesis for one of the best-established cases of functional specialization in the brain: the visual recognition of faces^2,6,7^.

Recent advances in deep convolutional neural networks (CNNs), which now achieve human level performance on some visual recognition tasks, allow us to test a prediction of our hypothesis^8–10^: If face recognition is functionally segregated in the brain because more domain-general visual representations simply do not suffice for this task, then the same should be true for any computational system, including CNNs. Prior work suggests that this may not necessarily be the case – CNNs optimized for generic object recognition transfer well to many other tasks^11–14^, including fine-grained discrimination within a category^12^. On the other hand, recent studies in other domains have found computational advantages of functional specialization in auditory processing^15^, and even spontaneous functional specialization for some high-level cognitive processes^16^. Thus it remains unknown, and not a priori obvious, whether the task of face recognition necessarily requires its own specialized machinery in any computational system, whether a brain or a machine. Here we address this question by measuring face and object recognition performance in CNNs trained to classify faces, objects, or both, testing whether high performance on both tasks happens only when distinct subsets of the network are allocated to each task. Note that in this work we do not address *spatial* segregation of function in particular locations in the brain, but the more general phenomenon of any *functional* segregation in which certain populations of units are more critical for one task than another, no matter where they reside spatially.

We first test object-trained and face-trained networks on both face and object recognition. One possible outcome is that CNNs trained only on object categorization will suffice for accurate face recognition, as they do for fine-grained discriminations within some other categories^12^. This finding would show that face recognition need not in principle require a specialized face system, instead favoring other accounts of the segregation observed in primate brains. However, we find that networks trained only on objects perform poorly on face recognition, and much worse than face-trained networks, supporting the computational account of why humans have specialized face systems. That finding raises a second question of what – if anything – must be built into a network for such a specialized face system to arise. One possibility, following long-standing evidence for innate domain-specific learning mechanisms in animals, is that face recognition could be learned from experience only if scaffolded upon built-in face-specific predispositions^17^ (such as an innate face template^18^). However, we find instead that face discrimination spontaneously segregates from object recognition in networks trained on both tasks, despite the lack of built-in face-specific inductive biases. This finding raises a third question of whether spontaneous task segregation in networks predicts the particular functional specializations observed in brains, or whether spontaneous task decomposition is a more pervasive property in networks. In fact, we find spontaneous segregation not only for faces, but also for other categories to varying degrees, revealing a general tendency for task segregation in networks, and opening the door to a wider investigation of the particular architectures, loss functions and training diets that determine *which* tasks will be segregated in networks, and by hypothesis also brains.

## Results

### Networks trained only on objects do not perform well on face recognition

To test whether object-trained CNNs suffice for face recognition and vice versa, we trained two randomly initialized VGG16 networks^19^, one on face identification (Face CNN; Fig. 1a in red) and one on object categorization (Object CNN; Fig. 1a in orange). We then asked how well the resulting feature spaces of each network trained on one task would support the other task, by decoding 100 held-out face and object categories based on the image representations (i.e., activation patterns) in the penultimate layer in each network (Fig. 1b). As expected, novel (i.e., untrained) face identities could be decoded accurately from the face-trained network (mean decoding accuracy: 82.2%) and novel objects could be decoded from the object-trained network (74.1%). However, the object-trained network performed significantly worse (29.3%) at face recognition than the face-trained network (p=3.72e-11, two-sided paired t-test), and vice versa for object recognition (Face CNN: 17.3%; p=1.82e-12, two-sided paired t-test). Thus, representations learned for one task do not readily transfer to the other—each task appears to benefit from specialized task-specific representations.

**Figure 1.**
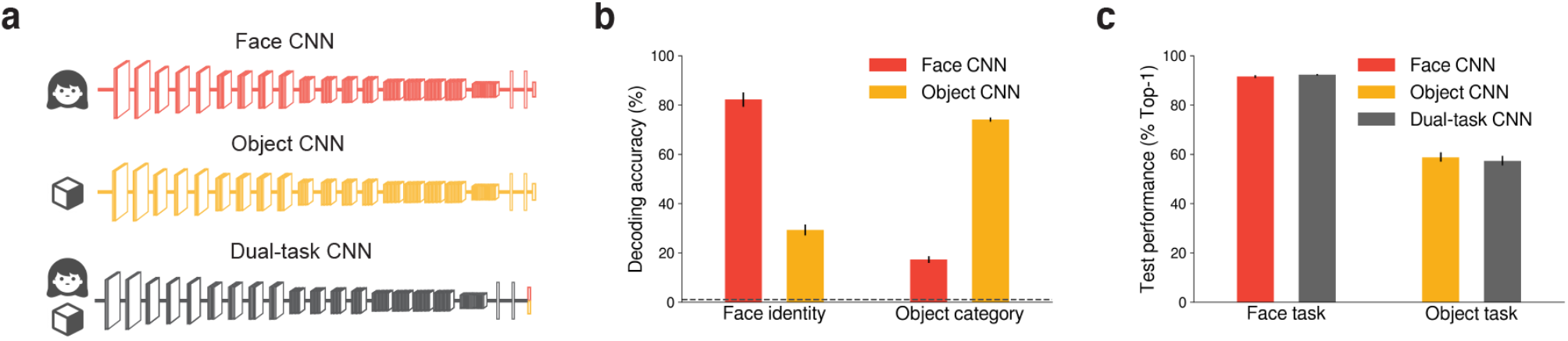
Distinct face and object representations in singly-trained CNNs, while a dual-task CNN performs well. **(a)** Three networks with VGG16 architecture were optimized, one on face identity categorization (Face CNN in red), one on object categorization (Object CNN in orange) and one on both tasks simultaneously (Dual-task CNN in gray). **(b)** Decoding accuracy of held-out face identities and held-out object categories using activation patterns extracted from the penultimate layer of the Face CNN and the Object CNN. The Face CNN outperforms the Object CNN in face decoding, and vice versa for object decoding. Thus, the representations optimized for each task do not naturally support the other. The dashed gray line indicates chance level (1%). Error bars indicate SEM across classification folds. **(C)** A dual-task CNN optimized on both tasks performed as well as the separate networks (% Top-1 accuracy on the test set). Error bars denote 95% CI bootstrapped across classes and stimuli.

### Spontaneous segregation for faces and objects in dual-trained networks

But might training a single network to perform both tasks discover a common high-performing feature space for faces and objects? To address this question, we trained a new network on both face identity and object categorization (Fig. 1a in gray)^15^. Surprisingly, this dual-task network performed nearly as well on each task as the network trained on that task alone (Fig. 1c). One interpretation of this result is that the network discovered a common feature space to solve both tasks, arguing against the hypothesis that specialization of function is required for high task performance. However, another possibility is that the network learned to segregate itself into two separate systems, one for face recognition and the other for object recognition, even though nothing was built into the network architecture or training regime to force it to do this.

To test this possibility, we performed a series of lesion experiments on the last convolutional layer (i.e., the final layer of feature extraction) of the model (Fig. 2a). First, we identified filters important for face recognition by measuring how ablating each filter (i.e., setting its output to zero) affected the loss for batches of face images (Fig. 2a, top panel) or object images (Fig. 2a, bottom panel) from the training set. We then ranked the filters according to their associated loss on each task. (Ranking by the selectivity of their response to faces or objects was less informative; see Supplemental Note 1.) Using a greedy procedure, we first selected and dropped the highest-ranking group (∼1.6%) of filters for each task, then selected the next highest-ranking group from the remaining filters in similar fashion but on novel batches of images. We repeated this process until there were no remaining filters left, resulting in all filters being ranked for their importance on each task (see Methods). Next, we tested the task specificity of this final ranking by lesioning the filters that most affected performance of one task (e.g. highest-ranking face filters) while measuring performance on the independent validation set for the face and the object task.

**Figure 2.**
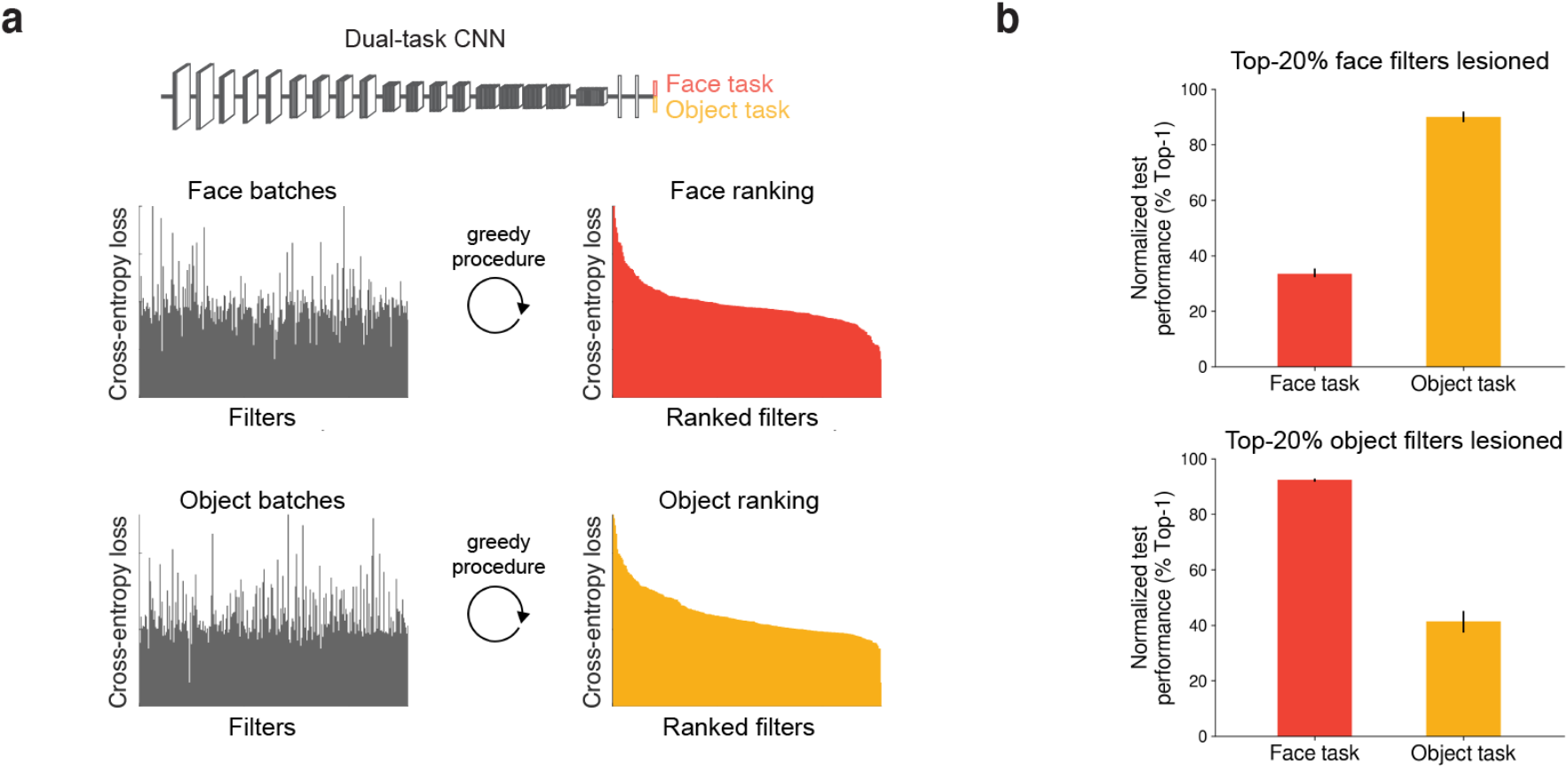
Lesion experiments in the last convolutional layer reveal spontaneous task segregation. **(a)** Schematic of lesion experiments for the last convolutional layer in VGG16. Each filter in the layer was ablated while measuring the loss to batches of face (top panel) and object (bottom panel) images. The filters were rank ordered by their corresponding losses to determine those that contribute most to face (red) or object recognition (orange). **(b)** Normalized performance of face and object tasks after lesioning the 20% highest-ranking filters for the face task (top) and the object task (bottom) in the last convolutional layer. Error bars denote 95% CIs bootstrapped across classes and stimuli.

This analysis revealed that lesioning the 20% highest-ranking face filters strongly impairs performance on the face task, but only minimally impairs performance on the object task, and vice versa for the highest-ranking object filters (Fig. 2b). These findings demonstrate a double dissociation in the network: face and object tasks rely on distinct features in the last convolutional layer. That is, the network spontaneously segregated itself into distinct subsystems for face and object recognition, despite the lack of any task-specific inductive bias that might have encouraged this outcome.

### Task segregation increases across layers, like the brain

How is this task segregation built up over layers of the network? In primate brains, processing of visual categories appears to share an initial set of common features in early stages of processing (retina, LGN, V1, V2, etc.), followed by branching into subsequent category-specific pathways (e.g., face, body, scene, etc.). Might dual-trained CNNs exhibit a similar organization? To test this possibility, we performed the same lesioning analysis in each convolutional layer individually (Fig. 3a). To quantify functional segregation, we defined a combined task segregation index. We defined task selectivity as the proportional drop on a given task (e.g. face task) when dropping any given group of filters together minus the proportional drop on the other task (e.g. object task) when the same group of filters are dropped, normalized by the sum of the two. We computed a face selectivity index and object selectivity index when the top-20% face-ranked and the top-20% object-ranked filters were dropped, respectively. The average of these two selectivity indices served as our combined task segregation index (see Methods). This segregation index is bounded between −1 and 1 where 1 indicates maximum segregation (e.g., lesioning filters would only impact the given task but not the other), 0 indicates no segregation (e.g., lesioning would impact both tasks equally) and −1 indicates inverse segregation (e.g., lesioning filters would only impact the other task but not the given task). An index of 1/3 indicates that the decrement in performance of the corresponding task is at least twice as large as of the other task.

**Figure 3.**
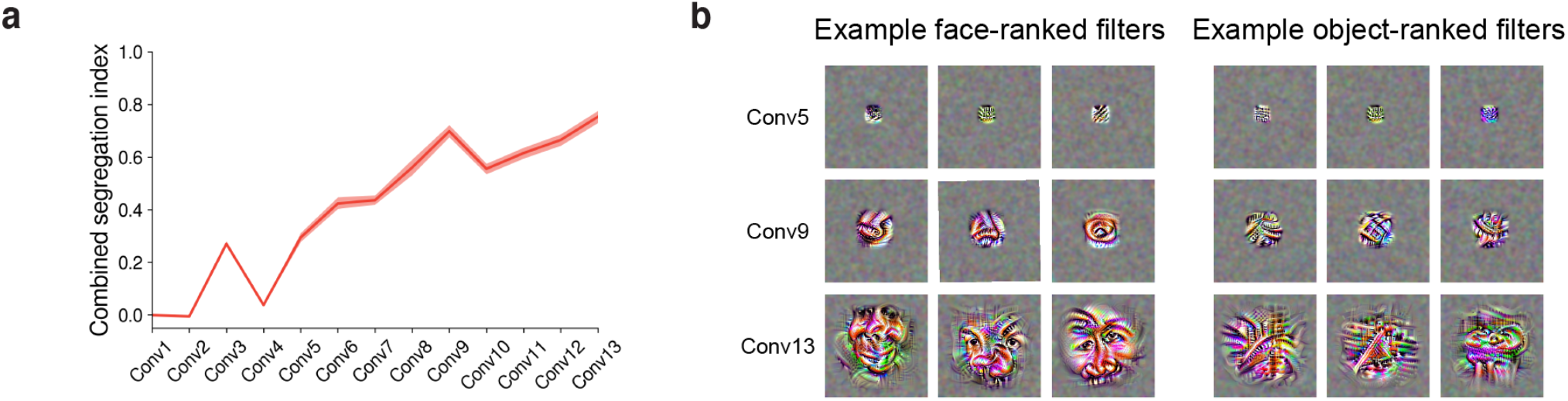
Spontaneous segregation of face and object tasks in mid-level processing stages. **(a)** Task segregation, measured as combined index of the differences in proportional drops in performance on the face and object task, when the 20% highest-contributing filters are dropped in each convolutional layer. Task segregation increased after the first convolutional layers to a maximum index of 0.75. Shaded area represents 95% CIs bootstrapped across classes and stimuli. **(b)** Images optimized to drive responses in 3 example filters among the top-10 selected filters for the face (left) and the object (right) task in convolutional layer 5, 9 and 13 (rows). The size of the receptive fields increases and features become more task-specific in later layers.

Indeed, we found that task segregation was small in early layers and increased with later layers, exceeding 1/3 for the first time at layer Conv6 (p=0, bootstrap test) and reaching 0.75 by Conv13 (see Fig. 3a). These results indicate that the processing of faces and objects gradually diverge at middle stages of processing in the network, and become highly segregated at later stages, much as we see in the primate brain.

The analyses above indicate that the network has effectively assigned some filters to the face task and other filters to the object task. But what features are these filters extracting? To find out, we visualized the preferred stimulus for each filter by presenting the network with a random noise input and modifying this input so as to maximize the activation of filters that ranked high on the face or the object task (Fig. 3b). While filters in early layers (e.g., Conv5) showed similar features across tasks, filters that were ranked high for the face tasks maximally responded to features that appear somewhat like face parts (e.g., nose, eyes) in mid-level layers (e.g., Conv9) and that appear to represent faces in a more holistic manner in late convolutional layers (e.g., Conv13). In contrast, object-specific filters were maximally activated by features and patterns that appear more generic, such as triangular shapes. These results show the development across the processing hierarchy of the distinctive features that each task relies on.

### Functional segregation does not arise for random tasks or distinct datasets for the same task

So far, we have shown that the double dissociation in primate brains between face and object recognition is recapitulated in CNNs optimized for both tasks, even though we did not build in any specific inductive bias to encourage the networks to discover this segregation. But perhaps the spontaneous segregation we found does not reflect task decomposition in any interesting sense, but would be found for any pairs of tasks, no matter how trivially they differ from each other. To test this possibility, we performed a control analysis to test whether a similar segregation would be found for random tasks. Here we randomly assigned 50% of the face and 50% of the object classes to a random task A, and the remaining face and object classes to task B (Fig. 4a). We then performed the same lesioning analysis based on these random tasks in the last convolutional layer (where we found strongest segregation for face and object tasks). We found no evidence of segregation based on these random tasks (Fig. 4b).

**Figure 4.**
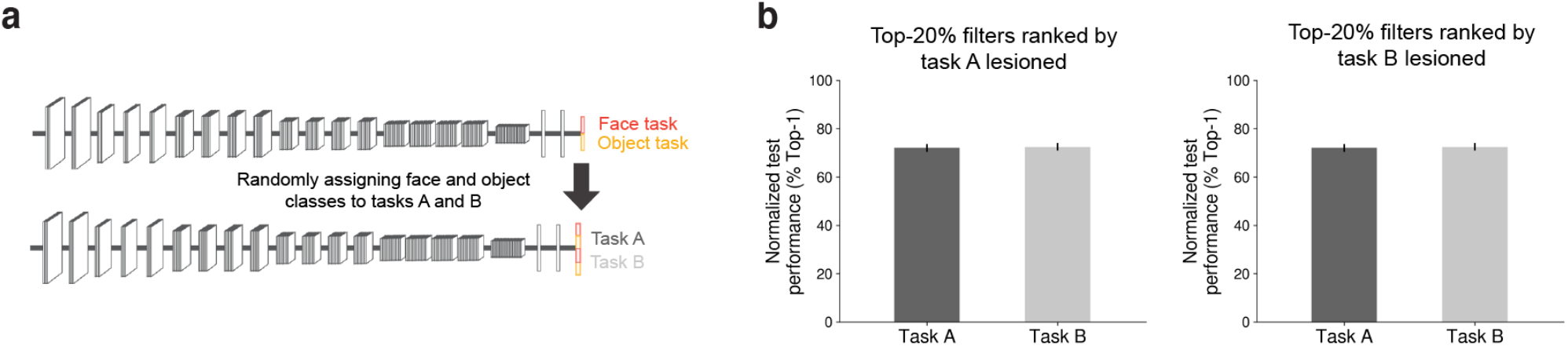
No segregation for random tasks. **(a)** Schematic of randomly assigning 50% of the face and 50% of the object classes to new tasks A and B. Each filter in the last convolutional layer was ablated while measuring the loss to batches of images belonging to task A or task B. Using a greedy procedure, the filters were rank ordered by their corresponding losses to determine those that contribute most to task A or task B. **(b)** Normalized performance of tasks A (dark gray) and B (light gray) after lesioning the 20% highest-contributing filters for task A (left) and task B (right) in the last convolutional layer. Performance decrement through lesioning was smaller than for the original tasks (Fig. 2b) and affected both tasks equally. Error bars denote 95% CIs bootstrapped across classes and stimuli.

The combined task segregation index was 0.01 (not significantly different from zero, p=0.596, bootstrap test), meaning that lesioning filters ranked highly on task A equally impacted performance on task A and task B, and vice versa. These results indicate that segregation does not develop for arbitrary tasks, but instead the network learns features that are specific to the face or object task, and that generalize across classes within – but not outside – each domain.

Could the functional segregation between tasks be explained by low-level differences or image collection biases in datasets? If low-level differences in tasks drive segregation, we would expect to find segregation starting in early layers of the network, where such low-level features are typically processed. However, in the layer-wise analysis of segregation above, segregation is minimal in the initial stages, only beginning to emerge at mid-level stages, suggesting that common low-level features are important for both tasks. Alternatively, each task might rely on distinct features, but impairment of these features in early layers might have less effect on performance. To distinguish between these two possibilities, we complemented this analysis with an analysis of how many of the top-20% ranked filters are shared between tasks (i.e., filters that ranked high on both tasks) in each layer. We found that the proportion of shared top-ranked filters across tasks was initially high, but strongly decreased with progressive layers (Supplementary Note 2), suggesting that the same filters contribute to both tasks in earlier layers, but each task relies on a distinct set of features in late layers. To further test whether low-level biases in datasets can drive segregation, we asked whether segregation can arise for the same task performed on two different image datasets. We found only a small degree of segregation in this case (Supplementary Note 3). Taken together, these control analyses suggest that the high degree of functional segregation we found for faces and objects is not due to simple dataset biases or low-level differences, but is instead driven by the distinct mid- to high-level visual features required for each task.

### Functionally segregated networks capture human behavior on faces and objects

Our findings of task segregation for faces and objects in dual-trained CNNs mirror the functional specialization observed in the human visual system. But are the learned feature spaces to perform both tasks also similar to the human visual system? Neural responses in task-specific areas for face and object processing have previously been linked to primate perceptual behavior in these tasks^4,20,21^. Are the learned representational spaces for faces and objects in the dual-trained network similar to those revealed in human behavior, and more so than the separate models optimized for only one task? To find out, we ran two behavioral experiments measuring the perceived similarity of pairs of face stimuli and pairs of object stimuli. For each task, we correlated the behavioral representational dissimilarity matrices (RDMs) of each subject for the corresponding stimuli with the RDMs obtained from each layer of the face-trained CNN (red), the object-trained CNN (yellow) and the dual-task CNN (gray) (Fig. 5). We found that the face-trained CNN was more similar to human face behavior than the object-trained CNN, and vice versa for object behavior, revealing a double dissociation. Importantly, each layer of the dual-task trained network matched behavior as well as or better than each of the singly-trained networks. These results show that the dual-task network captures human behavior in both face and object tasks, thereby supporting the conclusion that the learned solutions to perform both tasks resemble those in the human visual system.

**Figure 5.**
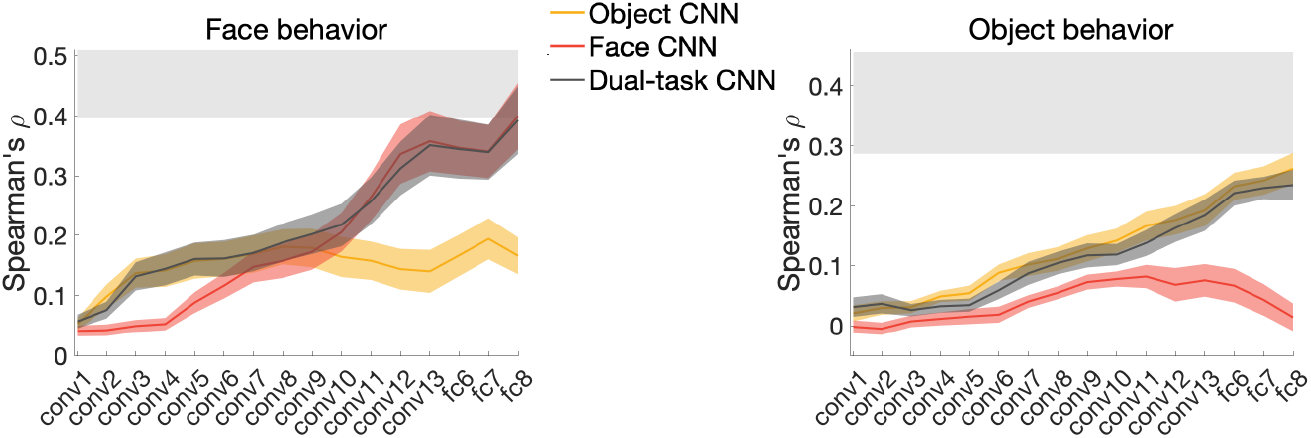
Dual-trained CNN is most correlated with behavior. Correlations between behavioral representational dissimilarity matrices (RDMs) for either face (left, n=14) or object (right, n=15) stimuli and layer-specific RDMs obtained from activation patterns in the Face CNN (red), the Object CNN (in yellow), and the dual-task CNN (in gray) to the corresponding stimuli. Color-shaded areas denote bootstrapped SEM across subjects. Gray-shaded horizontal bars indicate estimated noise ceiling based on the variability across subjects.

### Varying functional segregation for other visual categories

Thus, CNNs optimized for both face and object recognition, but with no domain-specific inductive biases, recapitulate both the double dissociation between face and object processing observed in the human brain, and the representational spaces for faces and objects revealed in human behavior. But are faces “special”, or might we also see spontaneous segregation of networks for the recognition of other categories (e.g., food, cars)? Note that we focus here on “natural” categories (i.e., those that frequently occur in the human visual diet) as we do not expect to find segregation in the human visual system for tasks that are not relevant to humans. One possibility is that only certain tasks are computationally distinct enough that they need their own separate processing mechanisms, and it is only these tasks that spontaneously segregate themselves in networks. Perhaps we could even “predict” which functions will be segregated in the cortex from the functions that spontaneously segregate themselves in networks. However, another possibility is that networks have a default tendency to segregate most natural tasks into distinct subnetworks. This might suggest that in brains too, functional segregation might be expected for most natural tasks that are important to us (or were important to our evolutionary ancestors). In that case our results would provide an explanation for why functional segregation is found in brains, but not for the particular functional specificities observed in the brain (e.g., for faces, places and bodies, but not for categories of similar evolutionary and modern-day relevance such as food^22^).

To test whether task segregation is found in networks even for natural tasks that have not been found to show segregation in the brain, we performed the same lesioning analysis on a dual-task network trained on visual food and object categorization (Fig. 6a). While food and objects showed weaker segregation than faces and objects in mid-level layers conv7 to conv9 (all p=0, bootstrap tests, fdr-corrected for number of layers), both networks showed similar degrees of segregation in the last convolutional layer (combined segregation index = 0.8 for food and objects; p=0.184, bootstrap test; fdr-corrected; Fig. 6b, green). Thus, spontaneous task segregation in networks predicts some specializations that have not been found in the brain. One potential explanation might be that food discrimination relies more on texture, while object categorization relies more on shape, and the “food” system is really a texture system in disguise^23^. Would another task that also requires fine-grained discrimination like face recognition, but that relies more on shape features, also show functional segregation? To test this idea, we trained a dual-task network on object recognition and car model/make discrimination (more fine-grained than most humans can perform). Interestingly, the car task showed later segregation than face and food tasks, exceeding a combined segregation index of 1/3 for the first time in layer conv9 (Fig. 6b, blue; p=0.013, bootstrap test, fdr-corrected). Moreover, while the degree of segregation in the last convolutional layer was still relatively high for objects and cars (max. combined segregation index = 0.56), it was significantly lower than for faces or food from layer conv3 onwards (all p=0, bootstrap test, fdr-corrected for number of layers and networks). Thus, CNNs show a widespread tendency to spontaneously segregate natural tasks to varying degrees, at least when half of their training is on that task, although their precise match to the brain needs to be further explored. It may turn out that all functional segregations found in the brain will also be found in CNNs, but the opposite may not be the case. Indeed, further investigation of the necessary conditions for spontaneous task segregation in artificial networks might reveal which factors determine the particular functional specificities observed in the brain^24^.

**Figure 6.**
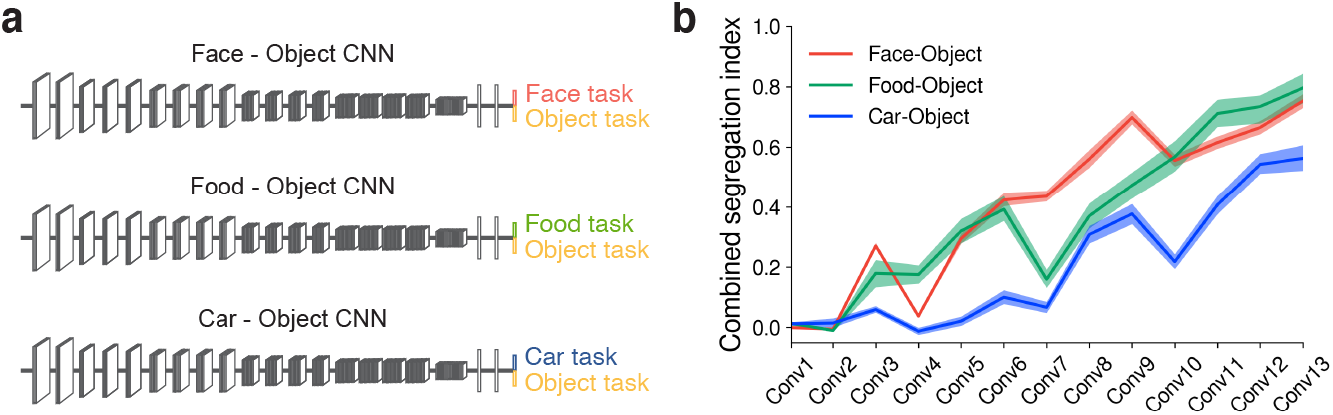
Spontaneous segregation to varying degrees for food or car recognition. **(a)** In addition to the dual-task model for face and object tasks (red), we trained one dual-task model on food (green) and object categorization, and another one on car (blue) and object categorization. **(b)** Task segregation was measured by lesioning the most-contributing filters for faces, food, and cars (respectively) and objects in each convolutional layer. Task segregation was found for all tasks to varying degrees. Task segregation for cars and objects increased later and to a lesser degree than for food or faces and objects. Color-shaded areas denote 95% CIs bootstrapped across classes and stimuli.

## Discussion

Our goal in this work was to understand why functional specialization is such a pervasive feature of brain organization, and to test the hypothesis that specialization in the brain may result from optimization (over development or evolution or both) for multiple natural tasks. This hypothesis predicts that even very different computational systems, with very different optimization procedures, may arrive at a similar solution^10^. We tested this prediction in CNNs for one of the best-established cases of functional specialization in the brain: face recognition. Supporting our hypothesis, we found that CNNs trained on object recognition perform poorly on face recognition and vice versa, and that CNNs trained on both tasks spontaneously segregate themselves into distinct subsystems for face and object recognition. We further showed that the dual-trained network also fits human perceptual behavior better than does either singly-trained network. And finally, although spontaneous network segregation is not found for trivially different tasks (like random tasks or the same task performed on two different data sets), it is found for other natural tasks including those for which brain specializations have not been reported. Taken together, these findings indicate that spontaneous task segregation is a widespread tendency in systems optimized to perform multiple natural tasks, lending support for the hypothesis that functional segregation in brains may reflect a computational solution given the structure of the tasks brains must solve.

Our findings also have implications for face perception in particular. Our results show that it is in principle not necessary to appeal to the social significance or meaning of faces to humans to understand how the face system develops or how it represents faces^25^. Our networks know nothing about why faces matter to humans; faces are just visual patterns to classify. And yet the networks “discover” the same functional dissociation between faces and objects that has long been reported in the human brain, and they “discover” a similar representational space for faces to that revealed in human behavior. These findings suggest that these properties of the human face perception system may result more from the computational structure of the task itself, than from the particular meaning faces have for humans. Of course, the networks used in our study could not achieve this performance on face recognition without extensive training on faces, which may also be true for minds^26,27^ and brains (but see^28–30^). To obtain this experience, networks would either need to be fed this information (as they were here), or if trained on more naturalistic data sets they may need some system to preferentially select faces for their own training input. For humans, faces indeed comprise a large percent of the perceptual input to human infants and adults^31–33^, and this has likely been true during much of human evolution. Indeed, it may be that the only inductive bias humans need to develop their face system is the already well-established early preference of infants to look at faces^34,35^.

Several limitations of this work should be noted. First, when we say a system has been *optimized*, that does not mean it is *optimal*. We have tested only a tiny subset of the possible architectures, data sets, and loss functions, and it remains possible that in some other corner of this large space a neural network could be found that will perform as well as the networks tested here and that may find a shared representational space for faces and objects. Thus we cannot argue that our results indicate that the human brain *must* segregate face and object processing to attain high performance on both – simply that under the circumstances we tested, we found segregation, resembling what we see in the brain. Second, the present work is agnostic on whether specialization in the brain is constructed through optimization over development or evolution or both. We trained CNNs to ask how they are functionally organized once optimized, not to ask how that optimization occurs. How this segregation arises in humans is bound to be very different from how it arises in CNNs trained with backpropagation. Third, this work does not address the spatial organization of functionally specialized systems in the brain, as the units and filters in the networks trained here have no analogue to location on the cortical sheet. These broader questions can be approached in future work using networks that explicitly model spatial topography^36,37^ and that are trained in a fashion more like human development^38^.

Finally, while our findings support a computational account of why functional specialization is found at all in brains, they do not yet accurately predict *which* mental functions should have specialized machinery in human brains. We find functional specialization for visual categorization of food in networks, but this has not been found in brains, despite the extensive interest and experience all of us (and our ancestors) have with visual discrimination of food. One possibility is that a visual specialization for food does in fact exist in brains, but at a finer spatial grain than current methods can detect. Another possibility is that the particular functional specializations found in CNNs might more closely match that found in brains for more brain-like architectures, training diets that more closely match human visual experience (e.g., our visual diet unlikely consists of 50% food), or different training regimes such as unsupervised or self-supervised loss functions^39–41^. A third possibility is that task-specific inductive biases need to be included in networks to match the particular specializations found in brains. Network optimization with and without task-specific inductive biases might help reveal which functional specializations rely on inductive biases in the brain^17^ and which do not.

Our findings dovetail with several prior lines of work. A few studies have shown performance advantages when branches are built into a network, with each branch trained on a different task, indicating an advantage of functional specialization^15,42,43^. Other studies have shown that even if each branch is not trained separately on a different task, the branches sometimes spontaneously differentiate themselves^40,44–46^. Separate functional responses can even arise in subsets of a network with no built-in branches^47,48^. Most impressively, one study found that the functional segregation that arose spontaneously in a network reflected distinct causal roles of each subnetwork in performance of different tasks^16^. Specifically, the authors in that study trained a RNN on twenty different cognitive tasks, and found that recurrent units developed into clusters, such that lesioning units in each cluster produced deficits in different subsets of the 20 tasks. Although it uses much simpler networks and tasks, this study mirrors our finding of spontaneous segregation of a network into distinct components, each causally engaged in a subset of the trained tasks. Lastly, several studies reported functional specialization for class-specific features using visualization techniques and lesioning methods^49,50^. While it has been proposed that class-specific units harm networks’ generalizability^51^, our control analysis using random tasks suggests that the emerging functional segregation is not class- but task-specific.

Perhaps the most exciting aspect of this work is the fact that we can move beyond simply describing the organization of the brain, to asking normative questions about why the brain is organized the way it is. We now not only have strong evidence that face processing is segregated from object processing in the brain, we have an understanding of why this might be the case: functional segregation is a natural consequence of optimization to solve multiple tasks. The stage is now set, and the methods are now in place, to tackle a suite of further why questions. Which visual categories require segregation and why these? What functional segregations are expected for domains other than visual classification? Should we expect to find functional segregation of rhythm and melody in music perception, or syntax and semantics in language understanding, based on the computational structure of these tasks? Network optimization and dissection provide powerful methods to tackle these^10^ and myriad other questions about why the brain is organized the way it is.

## Methods

### Task-optimizing CNNs on face and object recognition

To measure object categorization performance in a network trained on object categorization only (Object CNN), we trained a randomly initialized VGG16 network^19^ on 423 manually sampled categories of the ILSVRC-2012 database^52^. To avoid confounding the object task with scene or body categorization, for which segregation has also been found in the brain, we removed all categories from the original dataset that were scene-like (e.g., boathouse) or animals which include lots of bodies (and faces). We then manually chose categories that were prototypical objects (e.g., trumpet, hammer, coffee cup) from the remaining categories. For each of the 423 selected object categories, we used 1000 images for training and 200 for validation, for a total of 423,000 training and 84,600 validation images. We used similar training parameters as suggested in the original VGG paper^19^: stochastic gradient descent (SGD) with momentum with an initial learning rate of 10^−3^, a weight decay of 10^−4^ and momentum of 0.9. We manually reduced the learning rate twice to 10^−4^ and 10^−5^ when the training loss did not decrease for five epochs (i.e., full passes over the training set). To update the weights during training, we computed the cross entropy loss on random batches of 128 object images and backpropagated the loss. Each image was scaled to a minimum side length (height or width) of 256 pixels, normalized to a mean and standard deviation of 0.5, and data augmentation (i.e., 20% gray-scaled, randomly cropped to 224 x 224 pixel) was applied during training. The test images were scaled, normalized and center-cropped before extracting the classification. The resulting classification accuracy on the validation set provides a performance measure of a network that is free to learn task-optimized representations at all stages of visual object processing.

We measured face recognition performance achievable by the same architecture by training a VGG16 network on face identity categorization only (Face CNN). We trained the randomly initialized network on 1,714 identities (857 female) from the VGGFace2 database^53^. To allow for better comparison with humans, we tried to match the number of identities during training to the minimum number of people typically known by humans (range 1,000 – 10,000)^54^. We chose identities with a minimum of 300 images per identity and balanced female and male identities (857 each), otherwise the identities were randomly selected from the VGGFace2 database. To match the training set size to the Object CNN, we randomly chose 246 images per face identity for training, and 50 images per category for validation, for a total of 421,644 training and 85,700 validation images^15^. All other learning parameters were identical to the object network. The resulting classification accuracy on the validation set served as a measure for unconstrained face identity categorization performance.

### Decoding of within- and between-domain visual categories

To test whether representations and computations optimized for one task would transfer to the other task, we decoded exemplars of a held-out set of face identities and object categories from the activations extracted from both networks. We used 100 held-out face identities (50 female; 10 images per identity; 1000 images total) from the VGGFace2 dataset that were not included in the training set. For object decoding, we selected 100 categories from the THINGS database^55^ (10 images each; 1000 images total) that were prototypical object categories and did not overlap with the 423 categories the object CNN was trained on. We extracted the activation in the penultimate layer of each network (i.e., the last layer before the classification layer) to the 1000 face images and the 1000 object images, respectively. For each task and activations from each network, we trained and tested a 100-way linear support vector machine (with L2 regularization) on the corresponding activation patterns using a leave-one-image-out (i.e., 10-fold) cross-validation scheme.

### Training and testing dual-task networks

To determine whether representations could be learned to simultaneously support both tasks, we trained a dual-task CNN on face identity and object categorization (see Fig. 1a). We concatenated the face identity categories (1,714) and the object categories (423) to one task classification layer (2,137 categories in total) and used random batches of face and object images (batch size: 128) during training. All other learning parameters and the datasets were identical to the VGG16 training (see above). To measure the performance on the face and the object task, we computed the accuracy on the independent validation set separately for the face classes (1,714) and object (423) classes.

To test whether we would find segregation for other natural tasks, we additionally trained two dual-task networks: 1) We trained a network on object categorization and food discrimination using the Food101^56^ dataset. This dataset contains 101 food categories with 1000 images each. We used 900 images per category for training and 100 for validation for a total of 90,000 training and 10,000 validation images. Importantly, to maximize the distinction between the food and the object task, we removed all food classes from the object dataset (50 of the 243 original classes were food-related) prior to training. For the remaining 393 classes, we chose the number of training images for each class (231 images per class; 90,783 images in total) to match the number of the food images in the training dataset. 2) We trained a network on object recognition and fine-grained car model/make discrimination using the CompCars dataset^57^. To obtain enough images per class, we concatenated images from the same model/make but of different years into one class. In this fashion, we ended up with 1,109 classes with 45 images for training and 5 images for testing per class, for a total of 49,905 training and 5,545 validation images. We randomly chose 127 images from each class of the object dataset (i.e., all 423 classes for 53,271 images in total; none of these classes were car- or vehicle-related) to match the overall number of training images. Both of these networks were again trained with the identical training parameters as described for the model trained on faces and objects.

### Lesion experiments in dual-task networks

To test whether the dual-task network segregated the processing of faces and objects in the hidden convolutional layers, we performed lesion experiments^49^. First, for each convolutional layer, we identified putative task-specific filters by evaluating how much ablating that filter (i.e., setting its output to zero) affected the loss for 50 batches of face images and for 50 batches of object images, respectively, all taken from the training set. We then ranked the filters in each convolutional layer by how much they affected the loss on face images (face ranking) and the loss on object images (object ranking). Using a greedy procedure, we first selected and dropped the highest-ranking group (∼1.6%) of filters for each task, then selected the next highest-ranking group from the remaining filters in similar fashion but on novel batches of images (also taken from the training set). We repeated this process until there were no remaining filters left, resulting in all filters being ranked for the impairment they produced on each task when lesioned. For each layer, we then tested the task specificity of these filters by lesioning the top 20% filters that most affected performance according to the face or object ranking while measuring the performance on the independent full validation set for the face and the object task. To quantify functional segregation, we defined a combined task segregation index. First, we defined a task specificity (TS) index for a group of top-20% ranked filters *a* based on task A:

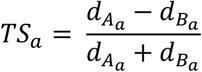

Where 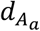 indicates the proportional drop on a given task A (e.g., face task) when dropping the top-20% filters *a* ranked by the corresponding task A altogether, 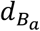 indicates the proportional drop on the other task B (e.g., object task) when the same group *a* of filters are dropped. We computed the task specificity index for the top-20% face-ranked and the top-20% object-ranked filters, respectively. The average of these two selectivity indices served as our combined task segregation index. This segregation index is bounded between −1 and 1 where 1 indicates maximum segregation (e.g., lesioning filters would only impact the given task but not the other), 0 indicates no segregation (e.g., lesioning would impact both tasks equally) and −1 indicates inverse segregation (e.g., lesioning filters would only impact the other task but not the given task). An index of 1/3 indicates that the decrement in performance of the corresponding task is at least twice as large as of the other task.

### Visualization of filters

To better understand the nature of the top face- and object-ranked filters, we generated images that strongly drove responses of single units in the network, optimizing the input image to maximize the unit’s responses. We initialized the input image with random noise, then used gradient ascent to maximize the units’ responses, with an additional L2 regularization over pixel values to encourage shrinkage of pixel values towards zero. Each unit visualized was chosen such that the center of its receptive field corresponded to the center of the image. This was done after observing that units from the same filter yielded similar synthesized images (but with receptive fields that were not centered at the center of the image). A small learning rate of 1e-3 worked best for maximizing the loss function. In addition, periodic (every 10 iterations) blurring and random jittering was applied to the pixel values^50^ to enhance visualization, presumably by discouraging getting caught in local minima. Note that units in filters from later layers have larger receptive field sizes than those at earlier layers. The visualizations demonstrate this property by optimizing over the receptive field size and leaving noise for image pixels on the borders.

### Testing segregation in random tasks

To test whether arbitrary tasks would lead to segregation in dual-task networks, we created two random tasks, each composed of half faces and half objects. To create two sets of face classes, face identities were chosen randomly until half of all identities were selected. This procedure resulted in two mutually exclusive, but exhaustive sets of identities, Faces-1 and Faces-2. The same procedure was repeated for object categories, resulting in two mutually exclusive and exhaustive sets of categories, Objects-1 and Objects-2, of equal size. To create two random tasks, we assigned the identities in Faces-1 and the categories in Objects-1 to Task-A and likewise Faces-2 and Objects-2 to Task-B. It follows that Task-A and Task-B are also mutually exclusive and of equal size where each comprise of half the face identities and object categories.

By removing filters that degrade the performance of Task-A as constructed above, we expect to find that Task-B is equally affected and vice-versa since presumably high impact features for either task are also of high impact for the other. We performed the same greedy lesioning analysis on the dual-task network trained on faces and objects based, as introduced above, but using the two random tasks (each comprising of half face identities and half object categories). The resulting proportional drop in each task and the combined segregation index then served as a baseline for a null specificity measure in dual-task networks.

### Human subjects

Behavioral data from 14 subjects (7 female; mean age 25.9, SD = 4.33) from a previously published study^58^ were used to perform the representational similarity analysis on face stimuli. As described previously, all subjects provided informed, written consent prior to the experiment and were compensated financially for their time.

Another set of 15 subjects (8 female; mean age: 28.9, SD = 8.2) were asked to perform the same task on object stimuli. All subjects provide informed, written consent prior to the experiment and were compensated financially for their time. The Massachusetts Institute of Technology (MIT) Committee on the Use of Humans as Experimental Subjects approved both experimental protocols (COUHES No 1606622600).

### Stimuli and behavioral representational dissimilarities

To find out whether humans and CNNs trained to recognition faces (Face CNN), objects (Object CNN) or both (dual-task CNN) represent faces and objects similarity, we performed representational similarity analysis. The experimental design to obtain the behavioral data has been explained in detail previously^58^, so here we just briefly summarize the stimuli and task. To obtain behavioral similarities for faces, Subjects (n=14) performed a multi-arrangement task^59^ using 80 gray-scale face stimuli. Stimuli consisted of 16 celebrities, which varied orthogonally in gender and age, such that half were female and half were male and half of them were young (below ∼35 years) and half were old (above ∼60 years). Another set of subjects (n=15) performed the same task but using manually selected images of objects. Stimuli consisted of five gray-scale images of each of eight prototypical object categories (chair, cheeseburger, dice, fork, guitar, headphones, car, kettle) for a total of 40 images. Subjects performed the multi-arrangement experiment online using the meadows platform (www.research-meadows.com) on their own computer. During the task, subjects were instructed to arrange different subsets of the images based on their perceived similarity (“similar images together, dissimilar images apart”) by dragging and dropping them in a circle. After the completion of the experiment, the pairwise squared on-screen distances between the arranged images was computed, thus representing a behavioral representational dissimilarity matrix (RDM). For each subject, we extracted the lower off-diagonal data from the behavioral RDM to obtain a vector of pairwise dissimilarities used for computing the correlations. We additionally computed the noise ceiling for the representational dissimilarities given the inconsistencies across subjects using a method described previously. Briefly, we estimated the upper bound of the noise ceiling as the mean correlation of each subject’s vector of perceived dissimilarities with the group mean (including the subject itself). In contrast, the lower bound was computed by taking the mean correlation of each subject with all other subjects.

### Representational similarity analysis between humans and CNNs

To obtain representational dissimilarities on faces and objects for the singly- and dual-trained CNNs, we presented the same stimuli (80 face images, 40 object images) as used for the human subjects to the CNNs. For each CNN, we extracted the activation patterns to each image separately for each of the 13 convolutional layers and the three fully connected layers and computed the correlation distance (1 – Pearson’s *r*) between each pair of activation patterns. This resulted in one RDM per layer for each of the three CNNs. To compute the similarity between the human RDMs and the RDMs obtained for the CNNs, we rank correlated each subject’s behavioral dissimilarities vector with the corresponding CNN dissimilarities vectors. The average rank correlation across subjects served as similarity measure between human subjects and CNNs. We further computed the bootstrapped confidence intervals by bootstrapping the subjects and computing the correlation with the CNN RDMs 10,000 times and computed the 95% CI of the resulting distribution.

### Significance Testing

We obtained bootstrapped 95% confidence intervals (CIs) for the accuracy of all networks by bootstrapping across classes and images 10,000 times. To obtain 95% CIs for the combined segregation indices, we bootstrapped across classes and images for the original and lesioned networks and computed the combined segregation indices for each bootstrap. Significance of comparison between the combined segregations indices and critical values (i.e., zero or 1/3), or between combined segregation indices of different networks was obtained by using direct bootstrap tests and FDR-correction.

## Data availability

The face stimuli used in the behavioral online experiment have been previously made available at https://osf.io/gk6f5/. The object stimuli used for the behavioral online experiment will be made available at https://osf.io/ upon acceptance. Behavioral data will be made available upon reasonable request. The source data underlying Figures 1–6 and Supplementary Figures 1–3 will be made available on OSF upon acceptance.

## Code availability

To reproduce the relevant analyses and figures, the Python scripts and functions will be made available on www.github.com upon acceptance. The computational models as well as the code to train computational models and extract activations will be made available upon reasonable request.

## Acknowledgements

We thank Jesus Crespo for support with training and analyzing computational models, and members of the Kanwisher lab for valuable comments and feedback on the manuscript. This work was supported by a Feodor-Lynen postdoctoral fellowship of the Humboldt Foundation to K.D., NIH grant Grant DP1HD091947 to N.K and National Science Foundation Science and Technology Center for Brains, Minds, and Machines.

## Author Contributions

K.D. collected and analyzed the data, J.M. provided additional scripts to train and test the computational models, KD., J.M, A.K and N.K. discussed the design and analyses of the manuscript and wrote the paper.

