## Supplementary Information for "Brain-like functional specialization emerges spontaneously in deep neural networks"

Dobs et al.

**Supplementary Note 1: Comparison of selecting filters causally versus based on response selectivity**

**Methods.** In our main analysis, we used a “causal” test to identify filters more important for face than object recognition and vice versa. But fMRI studies in humans identify “face areas” simply as the regions that show a higher response to faces than objects in localizer scans[1,2]. Would this latter selectivity method, applied to networks, reveal the same kind of filters we discovered using our lesioning method? Specifically, do filters with the highest impact on face performance (i.e., hence lower rank order number for the face loss) also show higher mean activation to face than object images and vice versa for object filters? To find out, we measured the activation of each filter in the last convolutional layer to an independent set of 100 face images (one image of each of 100 novel identities) and 100 object images (one image of each of 100 novel object categories), not included during training of the dual-task network. We extracted the activation to each of the 100 face and object images, respectively, by taking the l2 norm across the unit responses in each filter. We computed a standard selectivity index, measured as the difference in mean activation for images belonging to one category (e.g., face images) versus the other category (e.g., object images), divided by the sum of both. We then correlated the category-selectivity index of the 50% highest-contributing filters in the last convolutional layer (i.e., 256 filters) of the corresponding task (e.g., face task) with their rank order.

To further test whether the top-20% category-selective filters (e.g., face-selective) would selectively impair the corresponding task (e.g., face task) but not the other task (e.g., object task), we lesioned the top-20% category-selective filters simultaneously and measured the proportional drop in performance on each task.

**Results.** We found a significant negative correlation (Spearman’s r: -.40; $p<1e-10$; Suppl. Fig. 1A) between the rank of the top-50% highest contributing face filters (selected via lesioning) and their face selectivity based on activation, indicating that filters that have a large impact on face performance, also respond more to face than object images. We found a similar correlation between the rank of the top-50% highest contributing object filters and their object selectivity index (Spearman’s r: -.47; $p<1e-14$; Suppl. Fig. 1A). However, when we ranked filters by their selectivity based on activation and lesioned the 20% most category-selective filters, these filters impacted performance on the corresponding task less than “causally” selected filters (Suppl. Fig. 1B). We thus conclude that in the case of CNNs trained on face and object recognition, lesioning is a superior method for selecting task-specific filters compared to measuring differences in response activation.

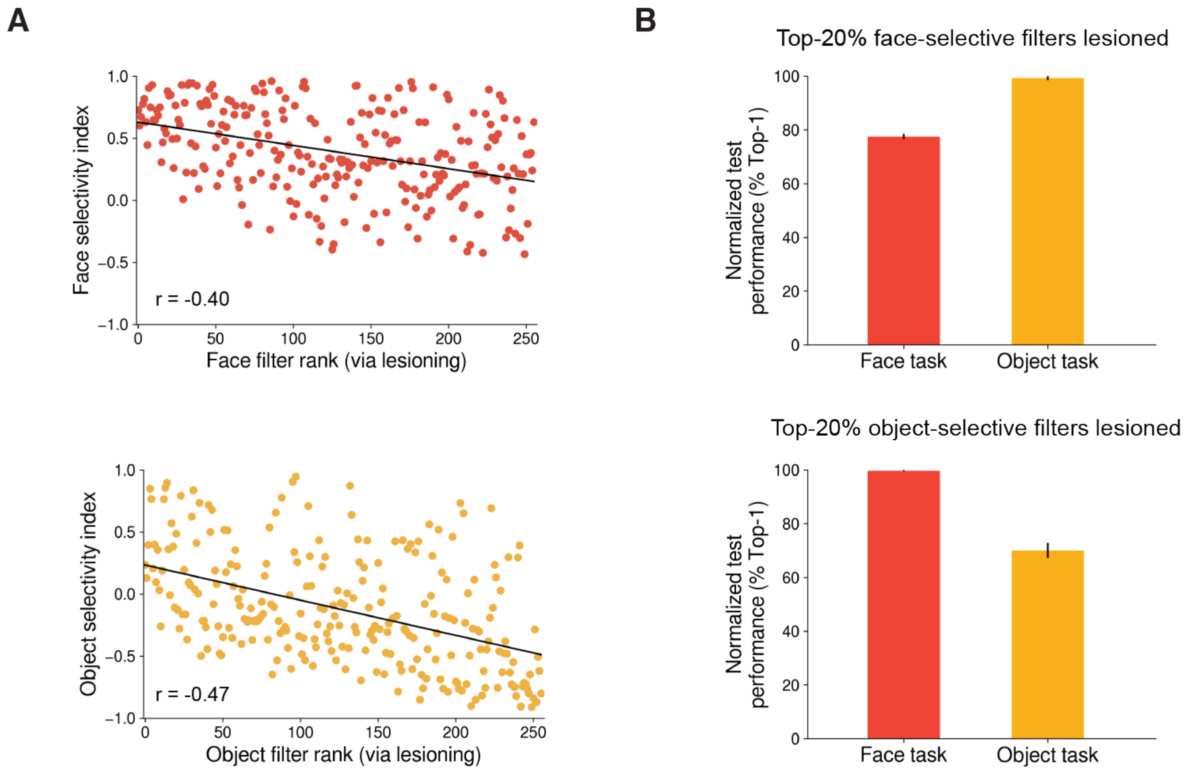


**Supplementary Figure 1 | Comparing category-selectivity based on activation to causally ranking filters. (A)** The rank of the top-50% highest contributing face filters (i.e., 256 filters) was significantly correlated with their face-selectivity index based on activation (r = -.40; top panel), suggesting that filters causally involved in the face task also responded more to face than to object images. Similar results were found for the top 50% object ranked filters (r = -0.47; bottom panel). **(B)** Lesioning the top-20% face-selective filters affected performance on the face task more than on the object task, but the drop in performance was lower than when selecting units based on lesioning (cf. Fig. 2). Similar results were obtained when lesioning object-selective filters. Error bars denote 95% CIs bootstrapped across classes and stimuli.

### Supplementary Note 2: Analysis of shared high-ranked units between tasks

**Methods.** To further test whether functional segregation for faces and objects is driven by mid- to high-level features, we performed an analysis of how many of the top-20% ranked filters are shared between tasks (i.e., filters that ranked high on both tasks) in each layer. If the low degree of segregation found in early layers of the network is indeed due to shared representations, the proportion of shared filters should be larger in early than in late layers.

**Results.** We found that the proportion of shared top-ranked filters across tasks was initially high, but decreased sharply with progressive layers (Supplementary Figure 2), suggesting that the same filters contribute to both tasks in earlier layers, but each task relies on a distinct set of features in late layers. This finding further supports the conclusion that functional segregation starts to emerge in mid-level layers of the networks, and thus is not likely being driven by low-level differences between tasks.

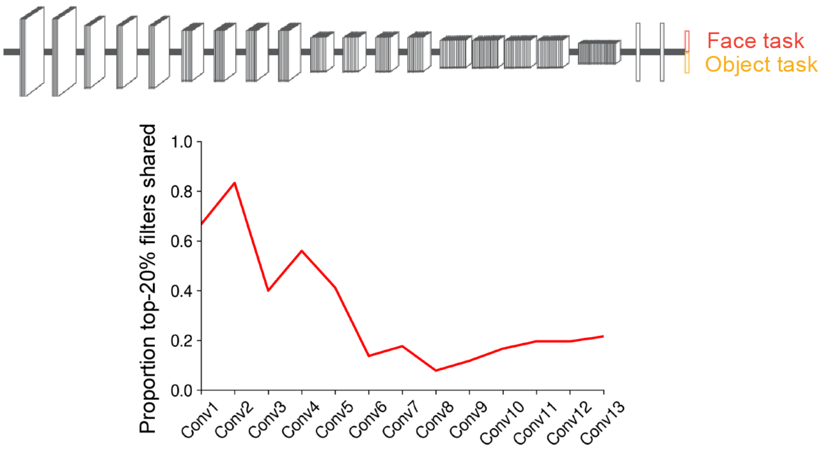


**Supplementary Figure 2 | Proportion of shared filters between face and object task decreased with progressive layers.** Analysis of the proportion of shared filters between the top-20% filters ranked by the face and the object task reveal a decreasing overlap with later layers of the dual-task network.

### Supplementary Note 3: Functional segregation for the same task based on different datasets

**Methods.** To further test whether biases in datasets can drive segregation, we asked whether segregation can arise for the same task performed on two different sets of images. Specifically, we trained a dual-task network on the face dataset used for the dual-task network trained on faces and objects (Face task 1, Suppl. Fig. 3 in red; 1,714 identity categories from the VGGFace2 dataset) together with 1,714 identities obtained from the CASIA-WebFace[3] (Face task 2, Suppl. Fig. 3 in purple) that were sampled on different selection criteria, cropping techniques, etc. We selected identities from the CASIA-WebFace dataset as follows. First, we ranked the identities by the number of images per identity (many of the identities contain only a few images per identity) and chose the 2000 identities with the largest number of images. Of these 2000 identities, we excluded all identities that overlapped with the VGGFace2 training set. From the remaining identities, we randomly chose 1714 identities with at least 63 images per class. Of these 63 images we used 58 for training and 5 for validation. To match the number of images between datasets, we also randomly selected 58 images per identity from the 1,714 identities of the VGGFace2 dataset. We trained the network with a 3428-way classification layer (1714 categories per dataset). We used the same training parameters and the same analysis to measure segregation in this dual-task network as we did for the dual-task network trained on faces and objects.

**Results.** We found that lesioning top-20% units in the last convolutional layer of the network impaired performance on both tasks much more similarly than what we found for faces and objects (Suppl. Fig. 3). The functional segregation was small in the last convolutional layer of the network (combined segregation index: 0.21) and significantly less than for faces and objects in the dual-trained network trained on faces and objects (combined segregation index: 0.75; p=0, bootstrap test). This finding further supports the conclusion that high degree of functional segregation we found for faces and objects is not due to simple dataset biases or low-level differences, but is instead driven by the distinct mid- to high-level visual features required for each task.

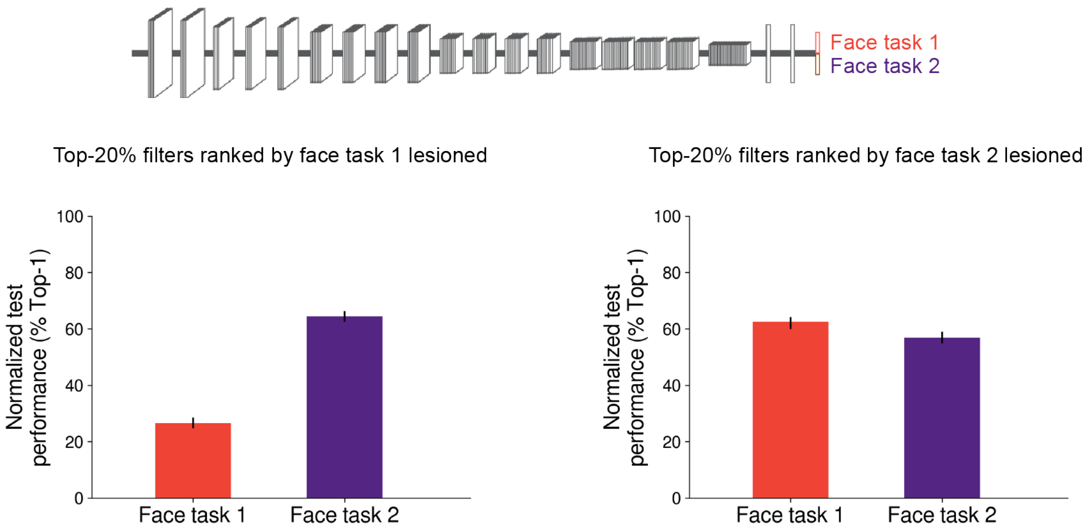


**Supplementary Figure 3 | Only small functional segregation for different datasets of the same task.** Normalized performance of face task 1 (red) and face task 2 (purple) after lesioning the 20% highest-contributing units for face task 1 (left) and face task 2 (right) in the last convolutional layer. Performance decrement through lesioning affected both tasks and to a similar degree. Error bars denote 95% CIs bootstrapped across classes and stimuli.
